# Monocyte-specific changes in gene expression implicate *LACTB2* and *PLIN2* in Alzheimer’s disease

**DOI:** 10.1101/2020.06.05.136275

**Authors:** Janet C. Harwood, Ganna Leonenko, Rebecca Sims, Valentina Escott-Price, Julie Williams, Peter Holmans

## Abstract

More than 50 genetic loci have been identified as being associated with Alzheimer’s disease (AD) from genome-wide association studies (GWAS) and many of these are involved in immune pathways and lipid metabolism. Therefore, we performed a transcriptome-wide association study (TWAS) of immune-relevant cells, to study the mis-regulation of genes implicated in AD. We used expression and genetic data from naive and induced CD14+ monocytes and two GWAS of AD to study genetically controlled gene expression in monocytes at different stages of differentiation and compared the results with those from TWAS of brain and blood. We identified nine genes with statistically independent TWAS signals, seven are known AD risk genes from GWAS: *BIN1, PTK2B, SPI1, MS4A4A, MS4A6E, APOE* and *PVR* and two, *LACTB2* and *PLIN2/ADRP*, are novel candidate genes for AD. Three genes, *SPI1, PLIN2* and *LACTB2*, are TWAS significant specifically in monocytes. LACTB2 is a mitochondrial endoribonuclease and PLIN2/ADRP associates with intracellular neutral lipid storage droplets (LSDs) which have been shown to play a role in the regulation of the immune response. Notably, *LACTB2* and *PLIN2* were not detected from GWAS alone.

## Introduction

Alzheimer’s disease accounts for 60-70% of the dementias, affecting 50 million people worldwide. Alzheimer’s disease can occur sporadically in the population or may be inherited. Early-onset AD (EOAD), that accounts for approximately 5% of AD cases, can be caused by autosomal dominant mutations that may be inherited or occur *de novo* in the *APP* (Goate *et al*., 1991), presenilin 1 (*PSEN1*) (Sherrington *et al*., 1995) or presenilin 2 (*PSEN2*) genes (Sherrington *et al*., 1996). However, most cases of AD are late onset (LOAD) and occur after the age of 65. Although one of the strongest risk factors for AD (both EOAD and LOAD) is being a carrier of the E4 allele of the *APOE* gene (Corder *et al*., 1993), recent GWAS have shown that variants in more than 50 loci are implicated in LOAD (Sims *et al*., 2020). Hence, it seems likely that multiple variants, each of small effect, acquired over a lifetime, may contribute to disease onset, defining LOAD as a polygenic trait (Escott-Price *et al*., 2015). These AD-risk genes are involved in multiple diverse biological pathways: such as the immune response, cholesterol metabolism, amyloid protein processing and APP metabolism (Jones *et al*., 2010; Kunkle *et al*., 2019; Sims *et al*., 2020) However, the mechanisms involved in the mis-regulation of gene expression that leads to AD have not been elucidated. Methods have been developed to test the association between changes in cell/tissue-specific gene expression and disease by predicting functional/molecular phenotypes into GWAS data (Gamazon *et al*., 2015; Gusev *et al*., 2016; Zhu *et al*., 2016; Barbeira *et al*., 2018) to decipher the functional relevance of disease - associated loci and to begin to define the mechanisms involved in their mis-regulation.

Innate immune activation in the brain is part of normal aging but it is also seen in AD pathogenesis. Up-regulation of innate immune activation genes is greater in normal aging than in AD, suggesting that inflammation is involved in the aetiology of AD at an early stage, before the detection of the disease (Cribbs *et al*., 2012). Microglia are the primary immune cells of the brain and have roles in innate immunity and in the development and maintenance of the central nervous system (CNS), maintaining neural circuits by synaptic pruning and by eliminating cellular debris. Microglia initiate an inflammatory response in the brain in response to amyloid-beta (Frautschy *et al*., 1998). However, as neuroinflammation inhibits neurogenesis (Monje *et al*., 2003) it is thought that chronic activation of microglia may contribute to cognitive decline and neuropathogenesis in AD. In addition, recent studies have implicated the microglia-mediated innate immune response in the development of Alzheimer’s disease (Sims *et al*., 2017; Tansey *et al*., 2018). Since peripheral monocytes differentiate into microglia-like macrophages within the CNS, we have utilised published expression and genetic data from monocytes at different stages of the inflammatory response (Fairfax *et al*., 2014) with the largest available Alzheimer’s disease GWAS summary statistics (Kunkle *et al*., 2019) to study genetically controlled gene expression in monocytes at different stages of differentiation. Given that the study of the correct tissue is important in identifying disease-relevant associations (Li *et al*., 2019), integrating monocyte expression with GWAS data using TWAS was expected to give insights into the genes within the immune response pathways that contribute to AD risk, along with the direction of effect and insight into the mechanisms involved the aetiology of AD.

## Materials and Methods

### Monocyte data

Monocyte expression data (Fairfax *et al*., 2014) was downloaded from array express (URLs). The corresponding genetic data was obtained from Dr Julian Knight University of Oxford under Licence (Fairfax *et al*., 2012, 2014, Naranbhai *et al*., 2015*b, a*)

### Monocyte expression data

Array Address Ids of the filtered probe set were extracted (15,421 probes) from the Fairfax *et al* Supplementary s file: 1246949stableS1.xlsx (URLs). An in-house script was used to convert Array Address Ids to Illumina Probe ids using the Illumina HumanHT-12_V4_0_R1_15002873_B array annotation information (URLs). Probes mapping to single genes for the 228 individuals with matched expression data available across all four monocyte cell strains were extracted (12469 probes, mapping to 9743 genes). Probes were mapped to GRCh37 (hg19) using the illumina_humanht_12_v4 chip annotation information using the Biomart package (URLs) and collapsed to genes using the ‘collapseRows’ function from the WGCA package (URLs) using the MaxMean method (Miller *et al*., 2011) in R.

### Monocyte genetic data

Genetic data files were aligned to the GRCh37 (hg19) using the Liftover tool (URLs). Plink 1.9 (URLs) was used for standard Quality-Control analysis (Anderson *et al*., 2010). Genetic data for the 228 individuals with matched expression data across all four cell states was extracted and SNPs were removed if their minor allele frequency (MAF) < 0.01, missingness of genotypes ≥ 0.02 or HWE < 10^−6^. A total of 625793 variants were retained for 228 people.

### AD summary statistics

International Genomics of Alzheimer’s Project (IGAP) is a large three-stage study based upon genome-wide association studies (GWAS) on individuals of European ancestry. In stage 1, IGAP used genotyped and imputed data on 11,480,632 single nucleotide polymorphisms (SNPs) to meta-analyse GWAS datasets consisting of 21,982 Alzheimer’s disease cases and 41,944 cognitively normal controls from four consortia: The Alzheimer Disease Genetics Consortium (ADGC); The European Alzheimer’s disease Initiative (EADI); The Cohorts for Heart and Aging Research in Genomic Epidemiology Consortium (CHARGE); and The Genetic and Environmental Risk in AD Consortium Genetic and Environmental Risk in AD/Defining Genetic, Polygenic and Environmental Risk for Alzheimer’s Disease Consortium (GERAD/PERADES). In stage 2, 11,632 SNPs were genotyped and tested for association in an independent set of 8,362 Alzheimer’s disease cases and 10,483 controls.

Meta-analysis of variants selected for analysis in stage 3A (n = 11,666) or stage 3B (n = 30,511) samples brought the final sample to 35,274 clinical and autopsy-documented Alzheimer’s disease cases and 59,163 controls. In the work presented here, GWAS Stage 1 summary statistics from Kunkle *et al* (Kunkle *et al*., 2019) were used. Common variants from this large and powerful study (21,982 cases and 41,944 controls) were filtered using the munge_sumstats.py (v2.7.13) script from the LD Score Regression (LDSC) software (URLs) and the hapmap 3.0 reference panel (URLs) resulting in 1,207,073 SNPs. Common variants from UK Biobank GWAS on parental AD : maternal and paternal Biobank data without IGAP (cases = 42,034, controls = 272,244) (Marioni *et al*., 2018) were filtered as described above, resulting in 1,188,072 SNPs.

### Expression weights

Expression weights for the monocyte data were computed using the R script : FUSION.compute_weights.R from the FUSION software (Gusev *et al*., 2016), using the binary plink files for the 228 individuals with matched expression data across all four cell strains and the corresponding expression data for each cell strain. Weights were packaged using the script from Oliver Pain (URLs). GTEx7 and CMC expression weights and the reference panel from the 1000 Genomes European population were downloaded from the FUSION web site (URLs).

UpSetR (URLs) was used to determine the number of genes computed in the expression weights in common and unique to each of the monocyte cell strains.

### TWAS

Expression weights were used in TWAS for autosomal chromosomes and excluding the MHC region, with the Alzheimer’s disease summary statistics using the R script FUSION.assoc_test.R from the FUSION software (Gusev *et al*., 2016). Results were corrected for multiple testing of multiple genes within each tissue using the Bonferroni method. Where a significant TWAS association was obtained from multiple genes in a locus (+/− 500 kb), conditional analysis was used to obtain statistically independent signals using the FUSION.postprocess.R script and default parameters.

## Results

### Transcriptome-wide association studies (TWAS)

Using the FUSION software (Gusev *et al*., 2016), we computed expression weights for naive and induced monocytes using published genetic and expression data from 228 healthy European individuals. Four monocyte strains were tested; naive CD14+, induced with lipopolysaccharide for 2 hours (LPS2) and 24 hours (LPS24) and with interferon-γ (IFN). For each strain, 1600-1800 genes were modelled with significant cis - heritable expression (Supplementary Table 1). The overlap in the number of genes for which cis-heritable expression was computed in each of the monocyte cell strains is shown UpSet plot (Supplementary Fig. 1)

Using the latest summary statistics for diagnosed AD (Kunkle *et al*., 2019) and the monocyte expression weights described above, we ran TWAS analysis on naive and induced monocytes (Supplementary Table 2) using FUSION software (Gusev *et al*., 2016). Across the monocyte cell strains, a significant TWAS association signal was generated for 13 genes (Fig.1 and Table 1). For these genes, we determined which of the monocyte cell strains had expression weights computed (Supplementary Table 3). Three of these genes: *STX3, LACTB2* and *PLIN2* had novel significant associations with AD. The remaining 10 genes (*APOE, BIN1, FNBP4, MS4A4A, MS4A6E, MYBPC3, PTK2B, PVR, SPI1* and *STX3*) have already been shown to be associated with AD in GWAS, giving confidence that we are detecting effects of genuine disease relevance using monocyte TWAS. Where there were multiple associated features at the same locus, we identified those that were conditionally independent using the post-processing script in the FUSION software. TWAS significant genes in the same region (those within a 100-kb window) were aggregated and conditional analysis on the other TWAS signals at the locus was undertaken for these regions (Supplementary Table 4 and conditional analysis plots). *STX3*, one of the novel TWAS significant genes was dropped after conditional analysis along with three of the genes already known to be associated with AD: *FNBP4, MYBPC3* and *MS4A6A*. Hence, we identified 9 genes with statistically independent TWAS signals (conditional p ≤ 0.05). *LACTB2* and *PLIN2* had novel significant associations with AD and *BIN1, PTK2B, SPI1, MS4A4A, MS4A6E, APOE* and *PVR* are already known to be associated with AD.

**Table 1.**
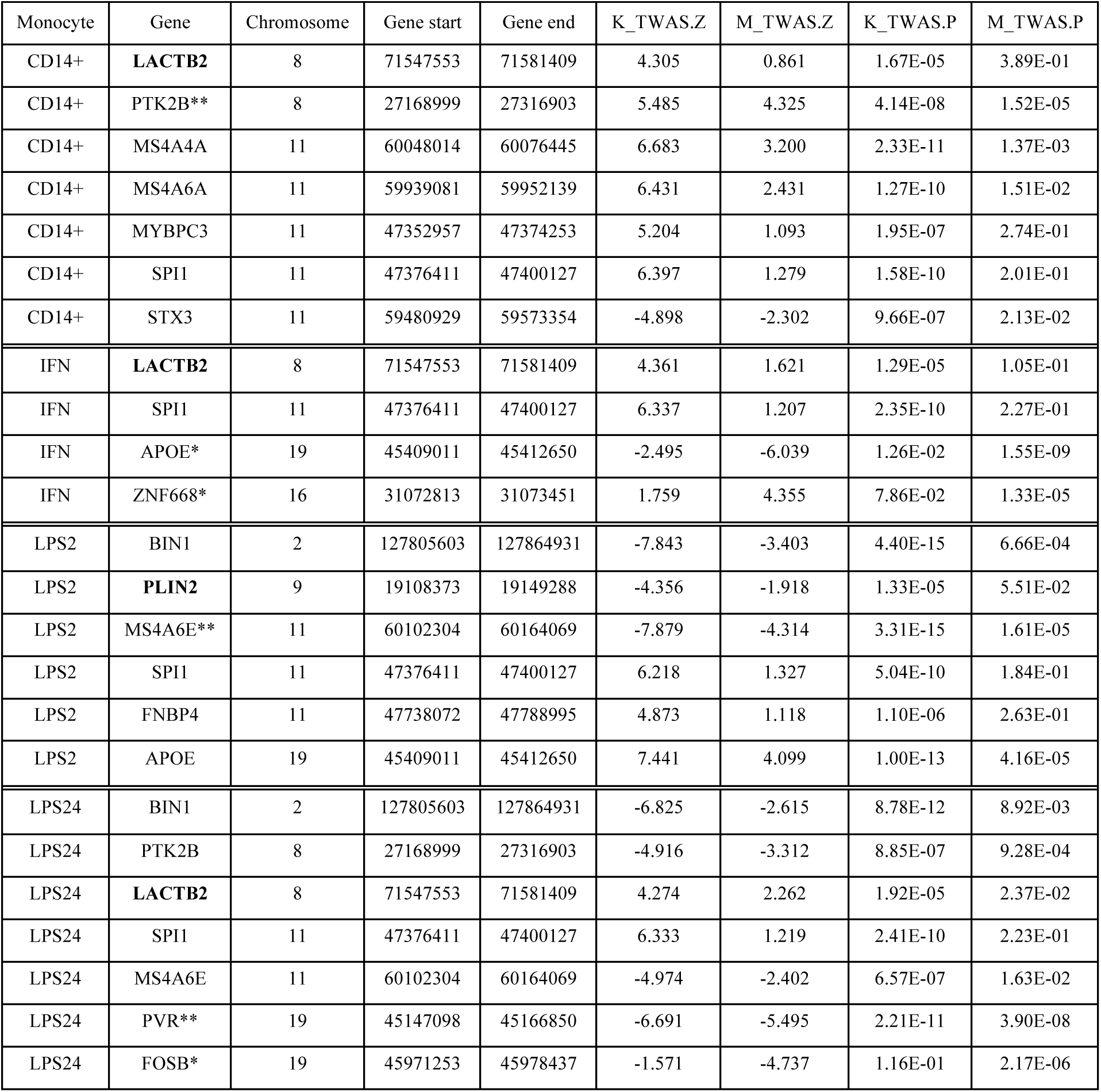
Transcriptome-wide association study (TWAS) test statistics for thirteen significantly associated genes for Alzheimer’s disease (AD) in monocytes. K_TWAS.Z and M_TWAS.Z denote the gene-level TWAS Z-score using the Kunkle 2019 (Kunkle *et al*., 2019) and Marioni summary statistics (Marioni *et al*., 2018) respectively. Genes without asterisks are TWAS significant using Kunkle summary statistics, * denotes TWAS significant genes using Marioni summary statistics, ** denotes TWAS significant genes in both Kunkle and Marioni TWAS. TWAS significant genes that are novel candidate genes for AD are shown in bold. TWAS_P values shown are not corrected for multiple testing, but all genes in the table are TWAS-significant in at least one of the TWAS after multiple testing correction (Bonferroni).

| Monocyte | Gene | Chromosome | Gene start | Gene end | K_TWAS.Z | M_TWAS.Z | K_TWAS.P | M_TWAS.P |
| --- | --- | --- | --- | --- | --- | --- | --- | --- |
| CD14+ | **LACTB2** | 8 | 71547553 | 71581409 | 4.305 | 0.861 | 1.67E-05 | 3.89E-01 |
| CD14+ | PTK2B** | 8 | 27168999 | 27316903 | 5.485 | 4.325 | 4.14E-08 | 1.52E-05 |
| CD14+ | MS4A4A | 11 | 60048014 | 60076445 | 6.683 | 3.200 | 2.33E-11 | 1.37E-03 |
| CD14+ | MS4A6A | 11 | 59939081 | 59952139 | 6.431 | 2.431 | 1.27E-10 | 1.51E-02 |
| CD14+ | MYBPC3 | 11 | 47352957 | 47374253 | 5.204 | 1.093 | 1.95E-07 | 2.74E-01 |
| CD14+ | SPI1 | 11 | 47376411 | 47400127 | 6.397 | 1.279 | 1.58E-10 | 2.01E-01 |
| CD14+ | STX3 | 11 | 59480929 | 59573354 | -4.898 | -2.302 | 9.66E-07 | 2.13E-02 |
| IFN | **LACTB2** | 8 | 71547553 | 71581409 | 4.361 | 1.621 | 1.29E-05 | 1.05E-01 |
| IFN | SPI1 | 11 | 47376411 | 47400127 | 6.337 | 1.207 | 2.35E-10 | 2.27E-01 |
| IFN | APOE* | 19 | 45409011 | 45412650 | -2.495 | -6.039 | 1.26E-02 | 1.55E-09 |
| IFN | ZNF668* | 16 | 31072813 | 31073451 | 1.759 | 4.355 | 7.86E-02 | 1.33E-05 |
| LPS2 | BIN1 | 2 | 127805603 | 127864931 | -7.843 | -3.403 | 4.40E-15 | 6.66E-04 |
| LPS2 | **PLIN2** | 9 | 19108373 | 19149288 | -4.356 | -1.918 | 1.33E-05 | 5.51E-02 |
| LPS2 | MS4A6E** | 11 | 60102304 | 60164069 | -7.879 | -4.314 | 3.31E-15 | 1.61E-05 |
| LPS2 | SPI1 | 11 | 47376411 | 47400127 | 6.218 | 1.327 | 5.04E-10 | 1.84E-01 |
| LPS2 | FNBP4 | 11 | 47738072 | 47788995 | 4.873 | 1.118 | 1.10E-06 | 2.63E-01 |
| LPS2 | APOE | 19 | 45409011 | 45412650 | 7.441 | 4.099 | 1.00E-13 | 4.16E-05 |
| LPS24 | BIN1 | 2 | 127805603 | 127864931 | -6.825 | -2.615 | 8.78E-12 | 8.92E-03 |
| LPS24 | PTK2B | 8 | 27168999 | 27316903 | -4.916 | -3.312 | 8.85E-07 | 9.28E-04 |
| LPS24 | **LACTB2** | 8 | 71547553 | 71581409 | 4.274 | 2.262 | 1.92E-05 | 2.37E-02 |
| LPS24 | SPI1 | 11 | 47376411 | 47400127 | 6.333 | 1.219 | 2.41E-10 | 2.23E-01 |
| LPS24 | MS4A6E | 11 | 60102304 | 60164069 | -4.974 | -2.402 | 6.57E-07 | 1.63E-02 |
| LPS24 | PVR** | 19 | 45147098 | 45166850 | -6.691 | -5.495 | 2.21E-11 | 3.90E-08 |
| LPS24 | FOSB* | 19 | 45971253 | 45978437 | -1.571 | -4.737 | 1.16E-01 | 2.17E-06 |

**Fig. 1.**
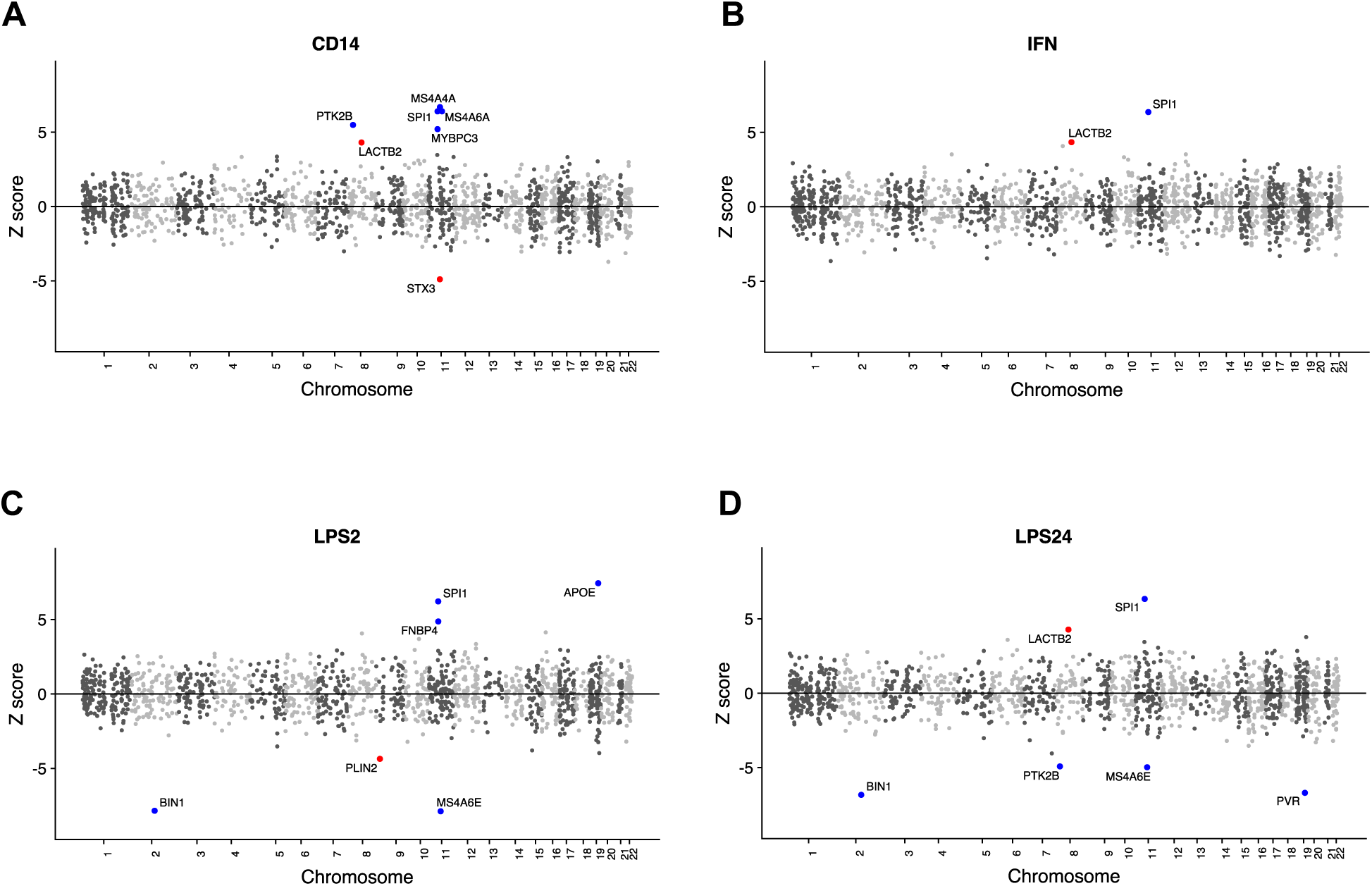
Mirrored manhattan plots of TWAS results. Genes are represented by coloured points plotted on the x axis by chromosome and genomic location. The y axis is the Z-score of the association between gene expression and Alzheimer’s disease in naive CD14+ cells (**A**), CD14+ cells induced with IFN (**B**), CD14+ cells induced with LPS for 2 hours (**C**) and 24 hours (**D**). Genes that show significant association after Bonferroni correction for multiple testing are shown with red (novel AD genes) and blue points (known AD genes) and named with black text.

### Comparison of TWAS from monocytes with TWAS from other tissues

To test whether AD-associated changes in gene expression were specific to monocytes, or whether they act across tissues, we performed TWAS analyses on the (Kunkle *et al*., 2019) GWAS summary statistics using published expression weights from the GTEx consortium (Lonsdale *et al*., 2013), the young Finns study, whole blood (YFS) (Raitakari *et al*., 2008), the Netherlands twin register peripheral blood (NTR), (Willemsen *et al*., 2010) and the CommonMind Consortium (URLs) (Supplementary Table 5). Correlation plots of the Z-scores from the TWAS analysis showed a high correlation between Z scores of the genes for which expression weights were computed in each of the monocyte cell strains (R > 0.75 in each case) (Supplementary Fig. 2). Correlation plots of the Z-scores from the TWAS analysis for each monocyte cell strain and GTEx whole blood (Fig. 2) show that *LACTB2* was TWAS-significant specifically in monocytes. In contrast, *PTK2B* was TWAS significant in CD14+ cells, LPS24 cells and whole blood (r=0.4-0.7, Fig. 2 and Supplementary Fig. 3). Similar correlation plots of the Z-scores from the TWAS analysis are shown for each of the monocytes cell strains versus, peripheral blood (NTR) (r=0.3-0.6, Supplementary Fig. 4) and CMC brain, dorsolateral prefrontal cortex (DLPFC) (r=0.3-0.4, Supplementary Fig. 5). We calculated the correlation co-efficients between all combinations of the AD relevant tissues, (GTEx and CMC Brain, GTEx and METSIM Adipose, GTEx and YFS whole blood, NTR peripheral blood and each of the monocyte cell strains (Supplementary Table 6). These results show that the largest correlation in Z-scores is between the monocyte cell strains, then between the monocyte cell strains and blood. The correlation between the monocyte cell strains and the brain tissues tested in all cases is low (r ≤ 0.5). The p value and number of genes in each pairwise correlation is summarised in Supplementary Table 5.

**Fig. 2.**
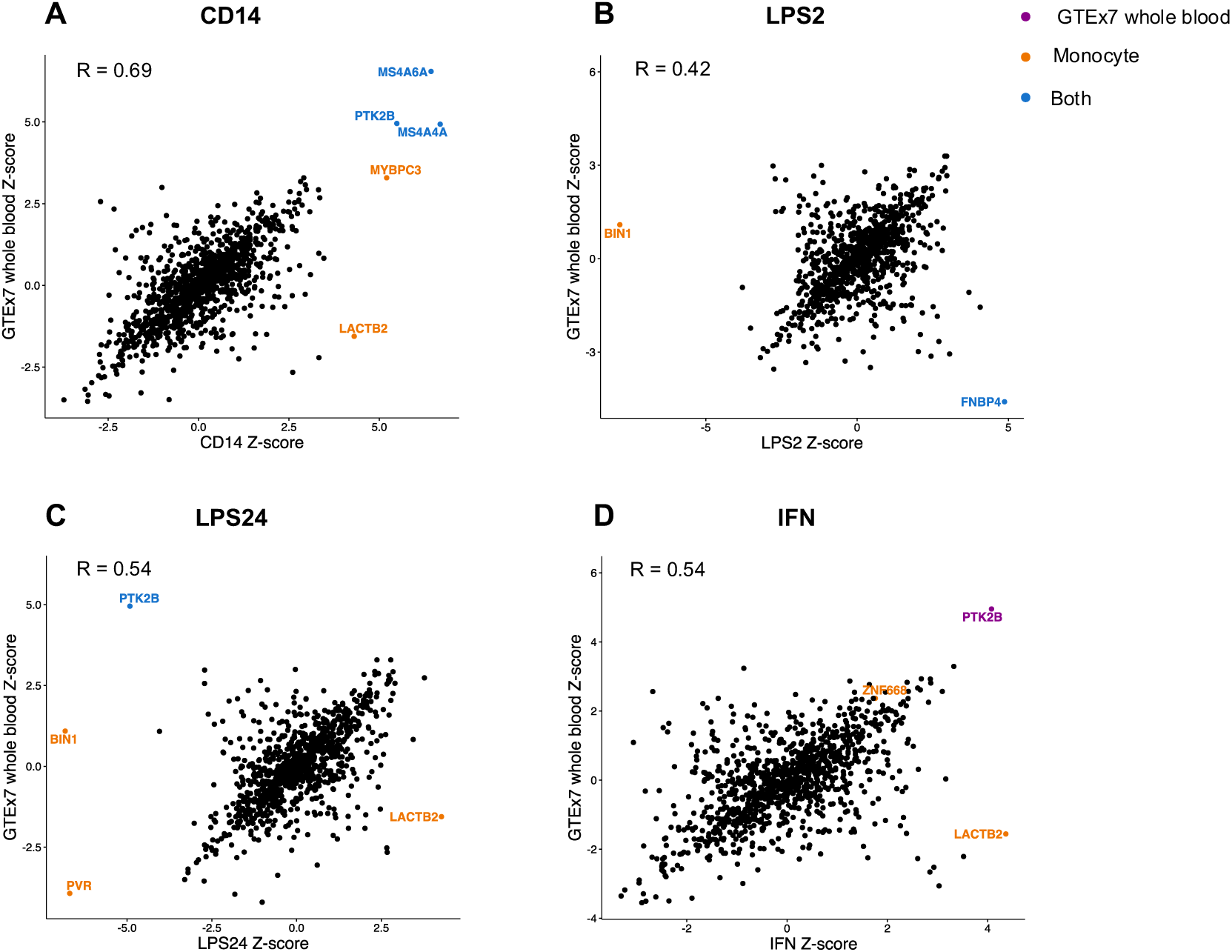
Correlation plots of Z-scores from TWAS in monocytes and whole blood. Genes are colour-coded according to their TWAS significance in cells in the monocyte lineage (orange), GTEx7 whole blood (purple) or both (blue). The increase in expression of LACTB2 associated with AD is only present in the monocyte lineage. R denotes the Pearson correlation co-efficient between the Z-scores in each case.

### Novel TWAS-significant genes, *LACTB2* and *PLIN2* are specific to monocytes

*LACTB2* is widely expressed (Uhlén *et al*., 2015). It was modelled with significant cis - heritable expression in 27 GTEx7 tissues which included whole blood, seven brain tissues, YFS blood and NTR blood. However, *LACTB2* was not TWAS-significant in any of these tissues (Supplementary Tables 5 and 7). *PLIN2* was modelled with significant cis – heritable expression in YFS blood and NTR blood, 2 GTEx7 cell lines, EBV-transformed lymphocytes and transformed fibroblasts, but not in any of the GTEx7 tissues. However, *PLIN2* was only TWAS-significant in LPS2-induced monocytes. This suggests that the association of the change in expression of both *LACTB2* and *PLIN2* with AD might be specific to monocytes.

### Functional analysis of the top eQTL variants in *LACTB2* and *PLIN2* in monocytes

The most significantly associated GWAS variant in *LACTB2*, rs7830986, detected in both naive and induced monocytes (LPS24 and IFN), was also the most significantly associated TWAS SNP but only in induced monocytes. rs7830986 is a synonymous variant in the *LACTB2* protein-coding gene and it is also within the lncRNA, *LACTB2-AS1* and an enhancer region. This SNP is likely to be involved in the regulation of *LACTB2* expression though the lncRNA or by affecting one of the transcription-factor binding sites in the enhancer region. The most significantly associated eQTL variant in naive CD14+ monocytes, rs13271014 is an intron variant in a regulatory region that overlaps the anti-sense lncRNA *LACTB2_AS1*.

The most significantly associated eQTL variant in *PLIN2* in LPS2 induced monocytes, rs10757004 is an intron variant in the *SAXO1* gene. This SNP is not in a regulatory region of the genome.

### TWAS-significant genes known to be in AD risk genes from GWAS

For the remaining TWAS-significant genes in monocytes that are known genome-wide significant AD risk loci (*APOE, BIN1, MS4A4A, MS4A6E, SPI1, PTK2B* and *PVR*) we summarise the results in the Supplementary Material.

### Comparison with TWAS from independent summary statistics

We replicated the TWAS using independent summary statistics from UK Biobank GWAS. This study used AD diagnosis based on a proxy by parental diagnosis design (Marioni *et al*., 2018). Overall, the correlation in gene-wide Z between the clinical AD (Kunkle *et al*., 2019) and AD-proxy analyses (Marioni *et al*., 2018) was modest (r∼0.2), although statistically significant (CD14: p= 2.2.×10^−16^, IFN: p = 4.9 × 10^−14^, LPS2: p=2.2× 10^−16^, LPS24: p=5.6×10^−16^) (Supplementary Fig. 6). Table 1 shows the Marioni TWAS Z-scores and p-values for the associated genes reaching statistical significance (after Bonferroni correction) in the Kunkle sample. It is notable that the sign of the Z-score which indicates the direction of effect, was identical in both samples for all genes. Of the 24 associations in Table 1, 15 achieved nominal significance (p<0.05) in the Marioni dataset. The known AD-risk genes: *PTK2B, MS4A6E* and *PVR* gave a significant p-value (after Bonferroni correction) in the Marioni dataset, however, *SPI1, MYBPC3, MS4A6E* and *FNBP4* did not. The novel AD genes, *PLIN2* and *LACTB2* showed minimal evidence for TWAS association in the Marioni dataset, although *LACTB2* showed a nominally significant association in LPS24. (Table 1, Supplementary Fig. 6).

## Discussion

Previously, CD14+ monocytes from a different dataset (BLUEPRINT) (Chen *et al*., 2016) have been used in a multi-tissue TWAS study of Alzheimer’s disease (Hu *et al*., 2019), alongside expression data from GTEx. However, analyses combining multiple tissues are best suited to detecting trait associations with genes highly expressed across several tissues, and may miss associations specific to a particular tissue (such as those with *SPI1*) (Li *et al*., 2019). Furthermore, the extent to which patterns of TWAS associations are consistent across monocyte cell strains and also brain and blood, has not been studied previously. Here, we found that the TWAS results from the four monocyte cell strains were strongly correlated with each other and they were also correlated with those from whole blood and peripheral blood. Correlation with CMC brain, (DLPFC) was less strong, a pattern also observed for the GTEx brain tissues.

We detected an association between changes in gene expression in both naive and induced CD14+ monocytes and Alzheimer’s disease. In all four monocyte cell strains, we detected a monocyte-specific association between an increase in the expression of *SPI1* and AD. The *SPI1* gene encodes PU.1, an ETS-domain transcription factor that activates gene expression during myeloid and B-lymphoid cell development (Scott *et al*., 1994) and in the CNS it controls the development of microglia. It is a master regulator that regulates several AD-risk genes (Huang *et al*., 2017). The most significantly associated eQTL SNP in our analysis was rs10838698. This SNP has been reported to be significantly associated with *SPI1* expression in monocytes and macrophages (Huang *et al*., 2017). Our analysis using AD summary statistics from Kunkle *et al* is in agreement with previous work and expands the association of a change in *SPI1* expression to both naive and induced monocytes. However, we note that the results for *SPI1* are not seen in TWAS using the AD summary statistics from Marioni *et al* (Marioni *et al*., 2018). This may be explained by the fact that these summary statistics were derived from UK biobank data using a proxy - phenotype design. The UK biobank data included family history information (parent or first-degree relative with AD or dementia) as a proxy-phenotype for the participants and this study is based on self-reporting of any type of dementia, which may contribute to ‘noise’ in the summary statistics. As a consequence, the effect sizes from the Marioni study are systematically smaller than those from the Kunkle study.

A significant TWAS association signal was detected for two novel risk genes for AD, *LACTB2* and *PLIN2*. This is interesting because in gene-wide analysis of the Kunkle *et al*. summary statistics using MAGMA (de Leeuw *et al*., 2015), *PLIN2* and *LACTB2* are not significant (p=1.28×10^−4^ for *LACTB2*, p=3.36×10^−2^ for *PLIN2*). The association between an increase in expression of *LACTB2* and AD was detected in both naive and induced monocytes. *LACTB2* (metallo-β-lactamase protein) is a mitochondrial endoribonuclease that cleaves ssRNA but not dsRNA or ssDNA. *LACTB2* is essential for mitochondrial function and cell viability and its overexpression in cultured cells causes a reduction in many mitochondrial transcripts (Levy *et al*., 2016). Mitochondrial dysfunction has been implicated in Alzheimer’s disease and across several other neurodegenerative disorders such as Parkinson’s and Huntington’s disease (Perez Ortiz and Swerdlow, 2019) and across neuropsychiatric disorders such as bipolar disorder, depression and Schizophrenia (Wu *et al*., 2019). Mitochondrial dysfunction results in defects in calcium signalling and apoptotic pathways, ATP-depletion which affects oxidative phosphorylation and in the depletion of mitochondrial DNA. Malfunction of mitochondrial processes implicated specifically in AD include oxidative damage (Nunomura *et al*., 2001) and accumulation of APP on mitochondrial membranes (Anandatheerthavarada *et al*., 2003). More recently it has been shown that mitochondria are important in immune cell regulation; influencing immune cell metabolism, differentiation, activation of the inflammatory response and regulating transcription (Angajala *et al*., 2018). Variants in mitochondrial DNA (mtDNA) have been associated with AD (Shoffner *et al*., 1993), but many of the processes that affect mitochondrial function, involve nuclear-encoded proteins (Ali *et al*., 2019), so it is more likely that variants in nuclear genes that affect mitochondrial function will affect AD pathology. However, it is also possible that mitochondrial dysfunction may itself lead to disease pathology. *LACTB2* may have role in regulating mitochondrial mRNA turnover, and since mitochondria regulate the immune response, defects in *LACTB2* gene function may affect the inflammatory response.

The association between a decrease in expression of *PLIN2* and AD was detected exclusively in LPS2 induced monocytes. After conditional analysis, the evidence that the association with AD is due to a change in gene expression is weak, however, this gene remains TWAS-significant and worthy of further investigation. The *PLIN2* gene encodes Perilipin-2 (Heid *et al*., 1998) which is one of the most abundant proteins in intracellular neutral lipid storage droplets (LSDs). LSDs contain a core of neutral lipids which are encapsulated in a monolayer of phospholipids and proteins. The lipids stored in LSDs are used in metabolism, membrane synthesis and cholesterol homeostasis and they are thought to be important in immune responses in myeloid cells (den Brok *et al*., 2018). LSDs are mobile within the cytoplasm and can move to associate with the endoplasmic reticulum (ER) and mitochondria. Many AD-risk genes are involved in lipid metabolism, for example *APOE* (Farrer *et al*., 1997), *ABCA7* (Karch *et al*., 2012), *CLU/APOJ* (Lambert *et al*., 2009) and *RORA* (Baker *et al*., 2019) and lipid metabolism pathways have been identified as key in AD (Jones *et al*., 2010). Thus *PLIN2* is a credible candidate AD-risk gene. Recently it has been shown that Perilipins are regulated by PPARγ which, when inhibited results in the down-regulation of PLIN2, affecting LSD formation (Tian *et al*., 2019). As RORA is a regulator of lipid homeostasis through PPARγ (Kim *et al*., 2017) we have connected two AD-risk genes involved in lipid metabolism through PPARγ.

## Conclusion

Using TWAS, we have shown an association between changes in gene expression in both naive and induced CD14+ monocytes and Alzheimer’s disease in known AD-risk genes and in two novel genes, *LACTB2* and *PLIN2*. The latter two novel associations were not detected through GWAS alone, highlighting the utility of TWAS approaches using a wide range of tissues. *LACTB2* and *PLIN2* are involved in mitochondrial function and lipid metabolism respectively and as such are likely to be credible candidate AD-risk genes. We have linked together two candidate AD-risk genes, *RORA* and *PLIN2* in the regulation of lipid metabolism through PPARγ, providing a more detailed insight into the lipid metabolism pathways involved in AD.

## Abbreviations

AD: Alzheimer’s disease
CD14+: Naive monocytes expressing CD14
CMC: CommonMind Consortium
DLPFC: Dorsolateral prefrontal cortex
EOAD: Early-onset AD
eQTL: Expression quantitative trait locus
GTEx: Genotype-Tissue expression
GWAS: Genome-wide association studies
HWE: Hardy Weinberg Equilibrium
IFN: CD14+ induced with interferon-gamma
LOAD: Late onset AD
LPS2: CD14+ cells induced with lipopolysaccharide for 2 hours
LPS24: CD14+ cells induced with lipopolysaccharide for 24 hours
LSDs: Intracellular neutral lipid storage droplets
MAF: Minor allele frequency
mtDNA: Mitochondrial DNA
NTR: Netherlands twin register peripheral blood
SNP: Single nucleotide polymorphism
TWAS: Transcriptome-wide association studies
YFS: Young Finns study, whole blood

## Acknowledgements

We thank the International Genomics of Alzheimer’s Project (IGAP) for providing summary results data for these analyses. The investigators within IGAP contributed to the design and implementation of IGAP and/or provided data but did not participate in analysis or writing of this report. IGAP was made possible by the generous participation of the control subjects, the patients, and their families. The i–Select chips was funded by the French National Foundation on Alzheimer’s disease and related disorders. EADI was supported by the LABEX (laboratory of excellence program investment for the future) DISTALZ grant, Inserm, Institut Pasteur de Lille, Université de Lille 2 and the Lille University Hospital. GERAD/PERADES was supported by the Medical Research Council (Grant n° 503480), Alzheimer’s Research UK (Grant n° 503176), the Wellcome Trust (Grant n° 082604/2/07/Z) and German Federal Ministry of Education and Research (BMBF): Competence Network Dementia (CND) grant n° 01GI0102, 01GI0711, 01GI0420. CHARGE was partly supported by the NIH/NIA grant R01 AG033193 and the NIA AG081220 and AGES contract N01–AG–12100, the NHLBI grant R01 HL105756, the Icelandic Heart Association, and the Erasmus Medical Center and Erasmus University. ADGC was supported by the NIH/NIA grants: U01 AG032984, U24 AG021886, U01 AG016976, and the Alzheimer’s Association grant ADGC–10–196728.

We thank Dr. Julian Knight (University of Oxford) for providing the genetic data from monocyte cells and Dr. Oliver Pain (King’s College, London), Dr. Matt Hill (Cardiff University) and Dr. Leon Hubbard (Cardiff University) for their computational help and advice in this work.

