## Supplementary Material for "Monocyte-specific changes in gene expression implicate *LACTB2* and *PLIN2* in Alzheimer’s disease"

#### TWAS-significant genes known to be in AD risk genes from GWAS

##### APOE

The *APOE* gene encodes apolipoprotein E which binds fat to form lipoproteins.
Its association with AD is well established through GWAS (Farrer *et al.*, 1997) and being a carrier of the E4 allele of the *APOE* gene is one of the strongest risk factors for AD (Corder *et al.*, 1993). We have shown that an increase in expression of *APOE* is associated with AD in LPS induced monocytes (2 hours) but this is not evident in monocytes induced with LPS for 24 hours. However, the same eQTL and direction of change expression is seen in GTEx lung tissue. The most significantly associated eQTL SNP, rs5157 is an intron variant of *APOC4* that is located in a regulatory region upstream of the *APOC2* gene.

##### BIN1

We have shown that a decrease in expression of *BIN1* is associated with AD in LPS induced monocytes (2 and 24 hours), but not in naive CD14+ or in IFN-induced monocytes.
The most significantly associated eQTL SNP, rs6710467 is an intergenic variant upstream of *BIN1* that has been associated with acute myeloid leukaemia (Lv *et al.*, 2017).
A decrease in expression of this gene is also associated with AD in ten of the GTEx7 tissues, using TWAS, but only two of these are AD-relevant; Brain Cerebellum and Brain Cerebellar Hemisphere. We note that previously published eQTL studies show similar results to our TWAS analysis for the association between a change in expression in *BIN1* and rs6710467 in monocytes (Fairfax *et al.*, 2012), whole blood and cerebellum (Kunkle *et al.*, 2019).

##### MS4A locus

The MS4A region is a genome-wide significant AD risk locus. Multiple genes in this locus are in high linkage disequilibrium (LD). After conditional analysis, we detected an association between an increase in expression of *MS4A4A* in naive CD14+ monocytes and AD. The association between and increase in expression of *MS4A4A* and AD has been shown in GTEx7 whole blood (rs573122) and thyroid (rs600550), (Supplementary Material Table 4 ) and (Kunkle *et al.*, 2019). In contrast, in the LPS induced monocytes (2 and 24 hours) we identified a statistically significant decrease in *MS4A4E* expression in AD. We did not detect any association between changes in expression of genes at this locus and AD in IFN-induced monocytes. Using TWAS, we did not detect an association between changes in expression of *MS4A6E* and AD in any of the GTEx tissues, YFS or NTR blood, which contradicts a previous report of an eQTL effect (an increase in expression) in whole blood (Zhernakova *et al.*, 2017). However, the decrease in gene expression associated with AD that we identified at the MS4A locus using TWAS may be specific to LPS - induced monocytes.

The most significantly associated eQTL SNP in the MS4A2 locus in naive monocytes was rs6591559 which has been reported previously as an eQTL in monocytes (Zeller *et al.*, 2010) This SNP is upstream of *MS4A4A* in the intergenic region between the *MS4A4A* and *MS4A4E* In LPS-induced monocytes the most significantly associated eQTL SNP is rs11824734, which was a downstream transcript variant of *MS4A4A*. Both of the SNPs in this locus are in regulatory regions, but they are not in a known transcription factor binding site.

**PTK2B**

We show that there is AD- associated increase in expression of *PTK2B* in CD14+ monocytes, whole blood and peripheral blood (GTEx7 and YFS respectively) whereas there is an AD-associated decrease in expression LPS-induced monocytes (24 hours). This association is not seen in other GTEx tissues, for example in Brain Cerebellum suggesting that the association of this gene with AD is specific to blood. In this TWAS analysis, the most significantly associated eQTL SNP, rs17057043 in CD14+ naive monocytes, peripheral (NTR) and whole (YFS) blood and in LPS-induced monocytes is located in intron 5 of *PTK2B*. Previously this SNP has been reported to affect the binding of IRF1, a transcription factor that regulates the innate and acquired immune response and it is involved in some of the cytokine responses to LPS (Rosenthal *et al.*, 2014). Interestingly, this eQTL shows cell-specific opposite directional effects between naive monocytes and LPS induced monocytes.

**PVR**

PVR /CD155 was first identified as the receptor for polio virus, but it has been shown to have many biological roles (Bowers *et al.*, 2017). It is important in the immune response, regulating cell-mediated immunity by promoting monocyte trans-endothelial migration (TEM) (Reymond *et al.*, 2004). *PVR* has been associated with AD in an AD-by-proxy meta-analysis (Marioni *et al.*, 2018). We have shown that a decrease in expression of *PVR* is associated with AD specifically in LPS - induced monocytes (24 hours), several GTEx tissues including subcutaneous adipose, but not in brain tissues or whole blood. The eQTL SNP, rs10426401 is in a regulatory region in the first intron of *PVR* and could affect several transcription-factor binding sites.

**SPI1**

In all four monocyte cell strains tested, we detected an association between an increase in expression of *SPI1* and AD. These associations were not seen in whole blood or peripheral blood, they are specific to monocytes. In naive CD14+ cells, IFN and LPS24 induced cells, the most significantly associated eQTL SNP was rs10838698. This SNP has been reported previously as a candidate variant affecting *SPI1* gene expression (Huang *et al.*, 2017). However, the intronic variant, rs755553 is the most significantly associated eQTL SNP in LPS2 cells in this analysis. This SNP is in a regulatory region of the *SLC39A13* gene.

### Supplementary Figures

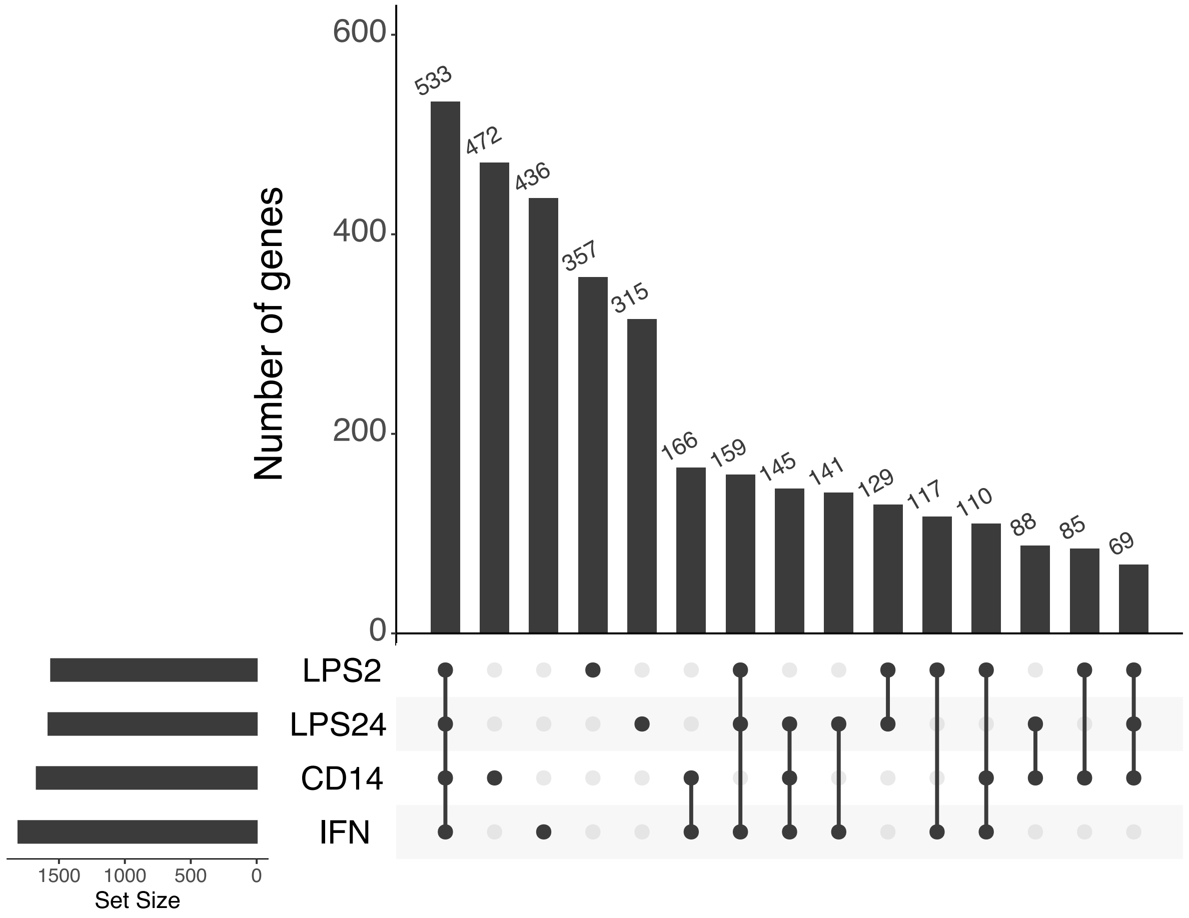


**Supplementary Fig 1. Genes for which cis-heritable expression was computed overlap between naive and induced monocyte cell types.** UpSet plots show the numbers of genes in common and unique to each of the monocyte cell types: Naive CD14+ cells (CD14), CD14+ cells induced with LPS for 2 hours (LPS2), CD14+ cells induced with LPS for 24 hours (LPS24) and IFN-induced CD14+ cells (IFN).

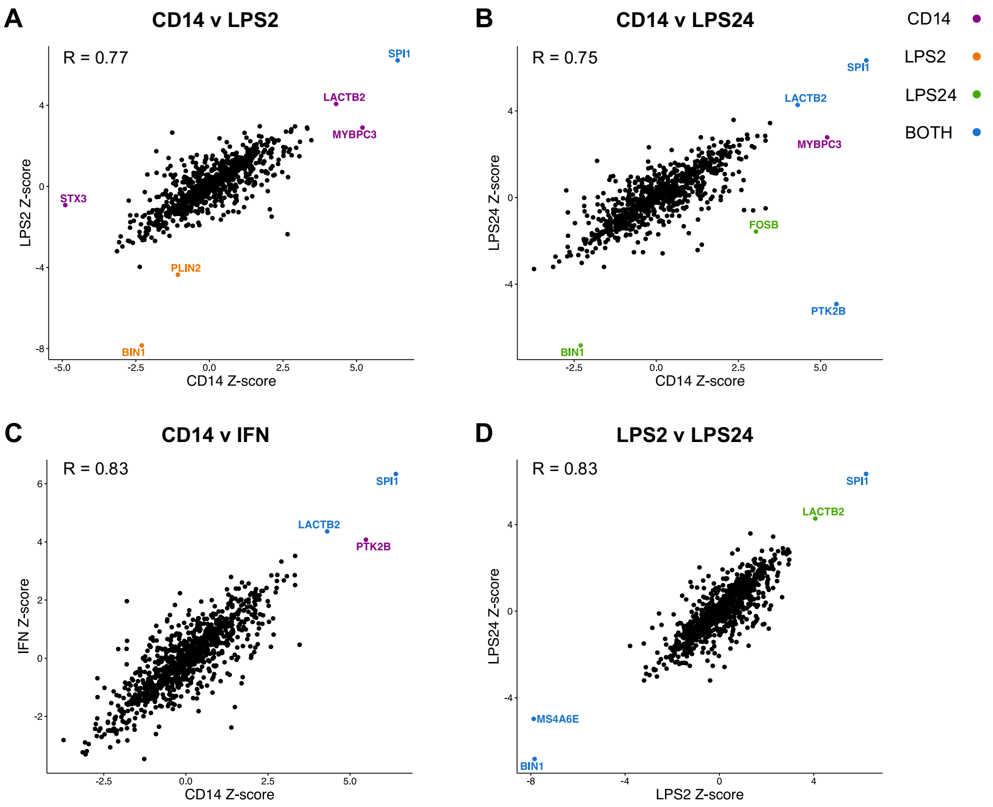


**Supplementary Fig 2. Correlation plots of Z-scores from TWAS in monocytes.** Genes are colour-coded according to their TWAS significance in cells each monocyte lineage: naive CD14 = purple, LPS2-induced =orange, LPS24_induced =green, or in both cell types (blue). The increase in expression of SPI1 associated with AD is present in all of the monocyte lineages and that of LACTB2 in naive CD14, LPS24 induced and IFN induced cells. R denotes the Pearson correlation co-efficient between the Z-scores in each case.

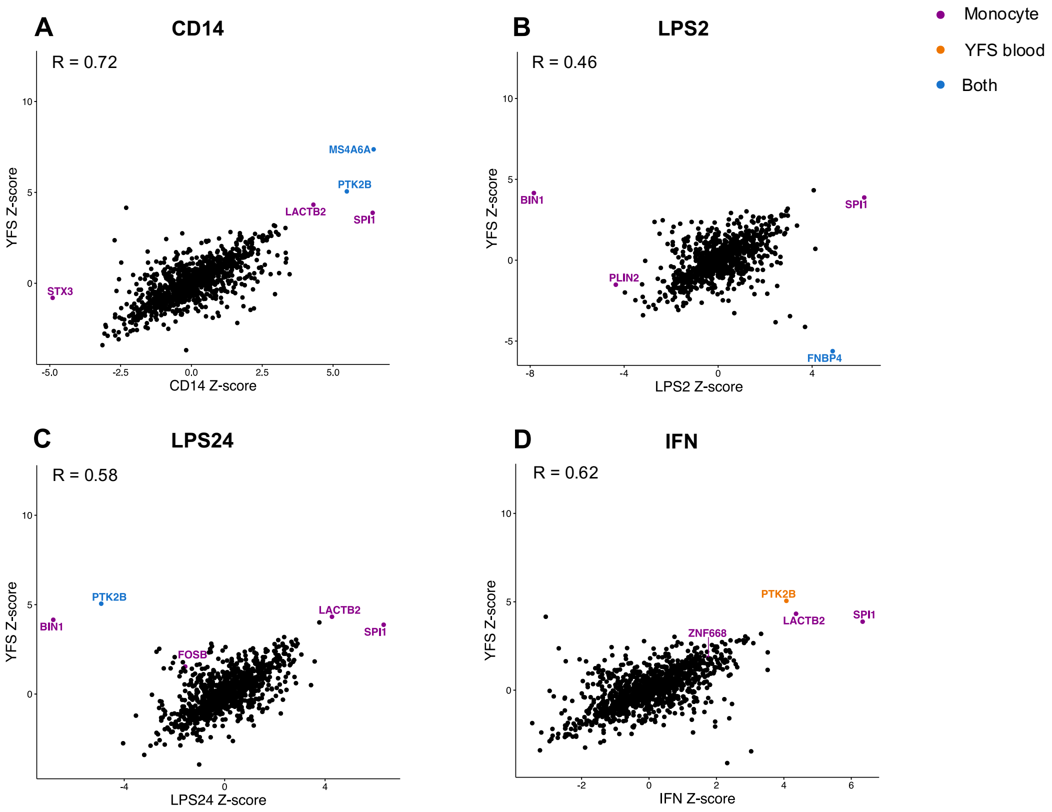


**Supplementary Fig 3. Correlation plots of Z-scores from TWAS in monocytes and YFS blood**. Genes are colour-coded according to their TWAS significance in monocyte lineages (purple), YFS blood (orange) or in both cell types (blue). R denotes the Pearson correlation co-efficient between the Z-scores in each case.

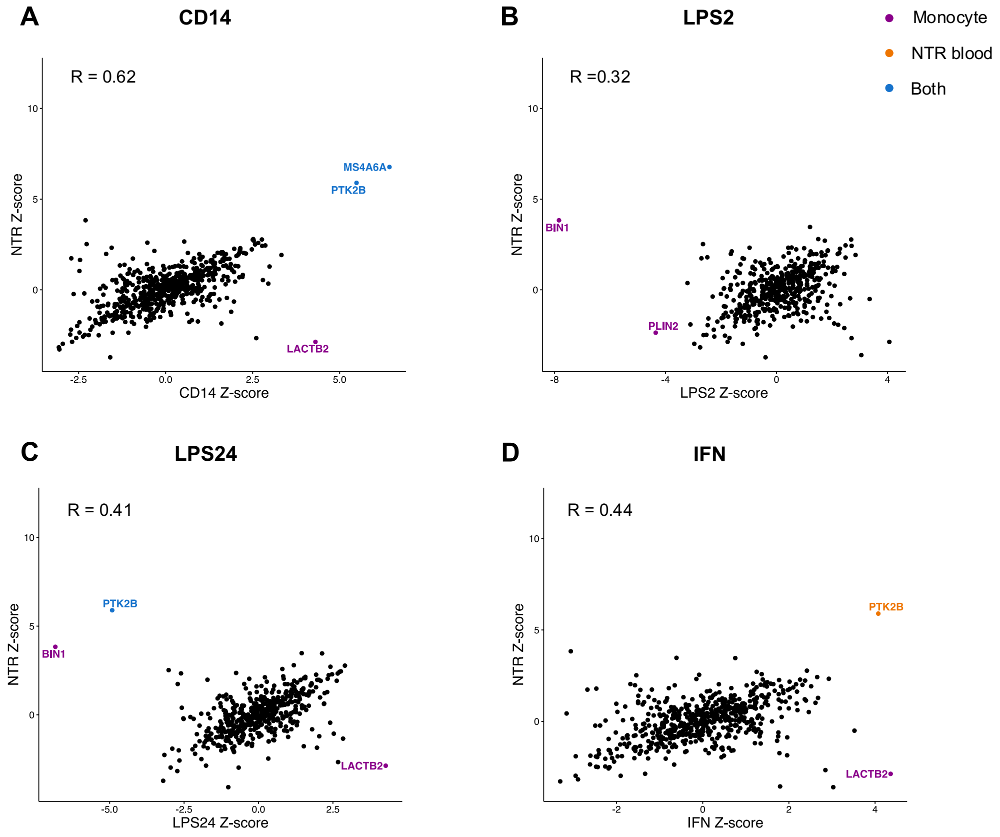


**Supplementary Fig 4. Correlation plots of Z-scores from TWAS in monocytes and NTR blood.** Genes are colour-coded according to their TWAS significance in monocytes (purple,) NTR blood (orange) or in both cell types (blue). R denotes the Pearson correlation co-efficient between the Z-scores in each case.

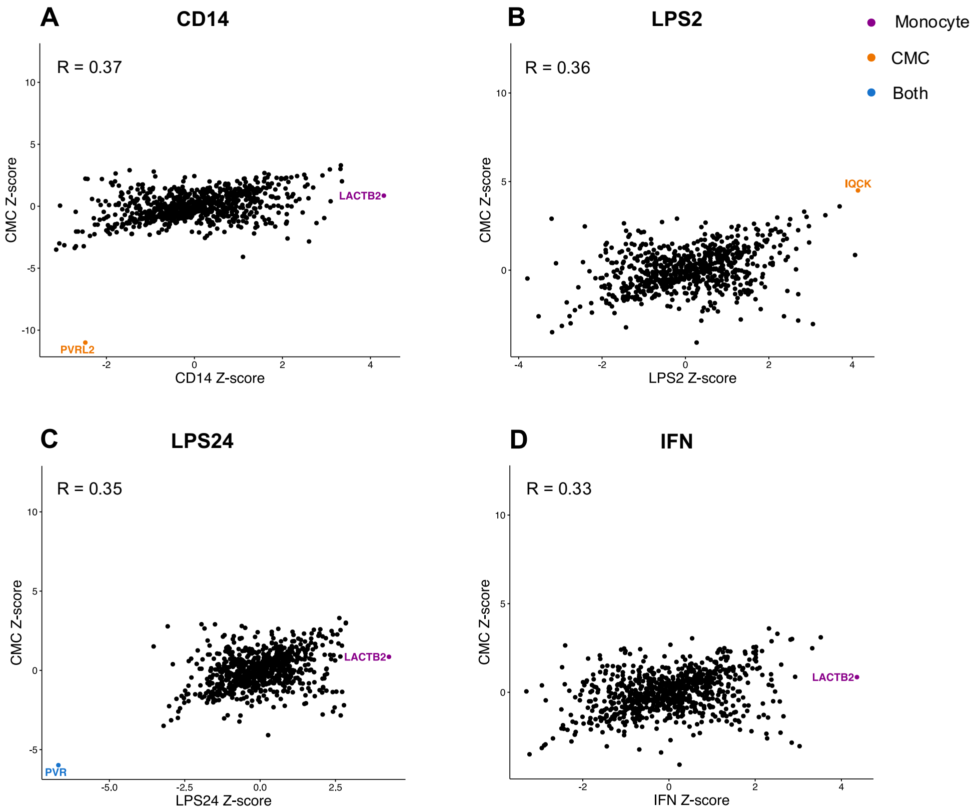


**Supplementary Fig 5. Correlation plots of Z-scores from TWAS in monocytes and CMC dorsolateral pre-fontal cortex (DLPFC).** Genes are colour-coded according to their TWAS significance in monocytes (purple), CMC - DLPFC (orange) or in both cell types (blue). R denotes the correlation co-efficient between the Z-scores in each case. R denotes the Pearson correlation co-efficient between the Z-scores in each case.

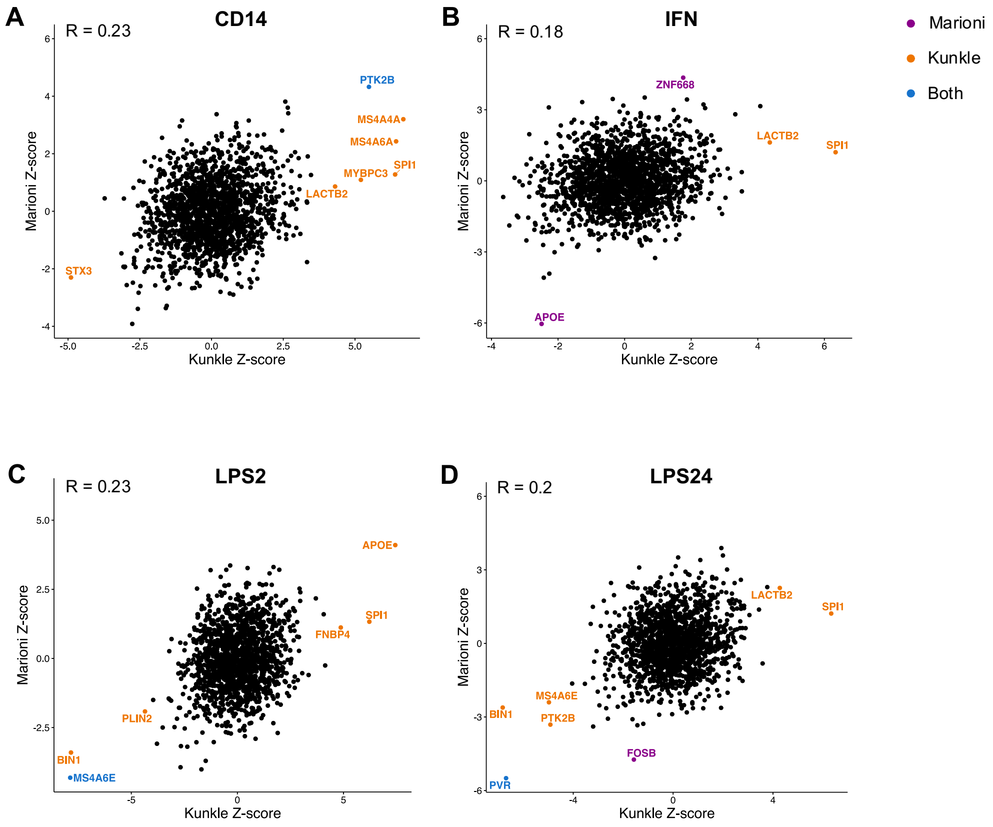


**Supplementary Fig 6**. **Correlation plots of Z-scores from TWAS using Kunkle and Marioni summary statistics.** TWAS significant genes (after Bonferroni correction) are colour-coded according to their being in the Kunkle (orange), Marioni (purple) or both (blue) summary statistics. The TWAS results for the PTK2B, MS4A6E and PVR genes are replicated in the Marioni TWAS in CD14+, LPS2 and LPS24 cells respectively. R denotes the Pearson correlation co-efficient between the Z-scores in each case.

### Supplementary Tables.

| Monocyte | No. of Genes with cis-heritable expression | No. of TWAS significant genes in Kunkle | No. of TWAS significant genes in Kunkle TWAS that survive conditional analysis | No. of TWAS significant genes replicated in Marioni * |
|---|---|---|---|---|
| CD14 | 1668 | 7 | 4 | 1 |
| LPS2 | 1559 | 6 | 5 | 1 |
| LPS24 | 1579 | 6 | 6 | 1 |
| IFN | 1807 | 2 | 2 | 0 |

No. of genes with significant TWAS association after Bonferroni correction
*Marioni summary statistics = UK Biobank GWAS on parental AD - meta analysis of log-odds and SEs from maternal and paternal AD (Marioni *et al.*, 2018)

**Supplementary Table 1. Summary of TWAS significant genes in the Kunkle and Marioni analysis for each monocyte cell type**. Genes with cis-heritable expression were used in the TWAS analysis for each cell type using Kunkle (Kunkle *et al.*, 2019) and Marioni (Marioni *et al.*, 2018) summary statistics. Three TWAS significant genes derived using the Kunkle summary statistics were replicated in the TWAS analysis using Marioni summary statistics: PTK2B, PVR, MS4A6E. The numbers of TWAS significant genes in each monocyte cell type obtained using the Kunkle summary statistics before and after conditional analysis are shown.

**Supplementary Table 2 (excel sheet ST2). Results of TWAS analysis from monocytes.** TWAS results using Kunkle summary statistics ((Kunkle et al., 2019)) and expression weights derived from monocytes: Naive CD14+ cells (CD14), CD14+ cells induced with LPS for 2 hours (LPS2), CD14+ cells induced with LPS for 24 hours (LPS24) and IFN-induced CD14+ cells (IFN).

| GENE/CELL TYPE | CD14 | LPS2 | LPS24 | IFN |
|---|---|---|---|---|
| APOE | | ● | | ○ |
| BIN1 | ○ | ● | ● | ○ |
| FNBP4 | ○ | ● | | |
| LACTB2 | ● | ● | ● | ● |
| MS4A4A | ● | | | |
| MS4A6A | ● | | | |
| MS4A6E | | ● | ● | |
| MYBPC3 | ● | ○ | ○ | |
| PLIN2 | ○ | ● | | ○ |
| PTK2B | ● | | ● | ○ |
| PVR | | | ● | |
| SPI1 | ● | ● | ● | ● |
| STX3 | ● | ○ | | |

**Supplementary Table 3. TWAS significant genes across the four monocyte cell types for which cis-heritable expression was computed.**

○ : denotes that a gene was present in the computed expression weights for a given cell type.

● : denotes that a gene was TWAS significant in the given cell type.

| GENE | Chromosome | Locus | Conditional analysis | Cell type | Known in AD GWAS |
|---|---|---|---|---|---|
| BIN1 | 2 | | | LPS2, LPS24 | (Seshadri *et al.*, 2010) |
| PTK2B | 8 | | | CD14, LPS24 | (Lambert *et al.*, 2013; Kunkle *et al.*, 2019) |
| LACTB2* | 8 | | | CD14, LPS24, IFN | |
| PLIN2* | 9 | | | LPS2 | |
| SPI1 | 11 | CELF1 | | CD14, LPS2, LPS24, IFN | (Huang *et al.*, 2017) |
| FNBP4 | 11 | CELF1 | dropped | LPS2 | (Karch *et al.*, 2016) |
| MYBPC3 | 11 | CELF1 | dropped | CD14 | (Huang *et al.*, 2017; Katsumata *et al.*, 2019) |
| MS4A4A | 11 | MS4A | | CD14 | (Naj *et al.*, 2011) |
| MS4A6E | 11 | MS4A | | LPS2, LPS24 | (Naj *et al.*, 2011; Karch *et al.*, 2016) |
| MS4A6A | 11 | MS4A | dropped | CD14 | (Hollingworth *et al.*, 2011; Jansen *et al.*, 2019) |
| STX3 | 11 | | dropped | CD14 | |
| APOE | 19 | APOE | | LPS2 | (Farrer *et al.*, 1997) |
| PVR | 19 | APOE | | LPS24 | (Marioni *et al.*, 2018) |

**Supplementary Table 4. Summary of the TWAS significant genes across the four monocyte cell types.** 13 genes were TWAS-significant across the four monocyte cell strains: Naive CD14+ cells (CD14), CD14+ cells induced with LPS for 2 hours (LPS2), CD14+ cells induced with LPS for 24 hours (LPS24) and IFN-induced CD14+ cells (IFN). After conditional analyses we identified 9 genes with statistically independent TWAS signals (shown in bold), two of these, LACTB2 and PLIN2 are novel candidate genes for AD (shown with *).

**Supplementary Table 5 (excel sheet ST5). Results of TWAS analysis from 52 tissues.** TWAS results using Kunkle summary statistics (Kunkle *et al.*, 2019) and expression weights derived from GTEx7 tissues, CMC. BRAIN (DLPFC), METSIM.ADIPOSE, YFS.BLOOD and NTR.BLOOD.

**Supplementary Table 6 (excel sheet ST6). Correlation of TWAS Z-scores between monocyte cell strains and AD-relevant tissues.** Correlation co-efficients (R2) and corresponding p-values for the correlation between the TWAS Z-scores using Kunkle summary statistics for each monocyte cell strain and relevant AD tissues: GTEx7 Brain, CMC_Brain (DLPFC) and GTEx7 whole Blood, NTR.BLOOD, YFS.BLOOD, GTEx7 Adipose and METSIM.ADIPOSE. the no. of genes in the overlap of the two variables is shown in each case.

| CELL_TYPE | GENE | CHR | GENE_START | GENE_END | K_TWAS.Z | K_TWAS.P |
|---|---|---|---|---|---|---|
| YFS.BLOOD | BIN1 | 2 | 127805603 | 127864931 | 4.15 | 3.27E-05 |
| NTR.BLOOD | BIN1 | 2 | 127805603 | 127864931 | 3.83 | 1.26E-04 |
| GTEx7_Whole_Blood | BIN1 | 2 | 127805603 | 127864931 | 1.09 | 2.76E-01 |
| **YFS.BLOOD** | **FNBP4** | **11** | **47738072** | **47788995** | **-5.62** | **1.88E-08** |
| **GTEx7_Whole_Blood** | **FNBP4** | **11** | **47738072** | **47788995** | **-4.60** | **4.31E-06** |
| YFS.BLOOD | LACTB2 | 8 | 71547553 | 71581409 | 4.32 | 1.53E-05 |
| NTR.BLOOD | LACTB2 | 8 | 71547553 | 71581409 | -2.87 | 4.12E-03 |
| GTEx7_Whole_Blood | LACTB2 | 8 | 71547553 | 71581409 | -1.56 | 1.19E-01 |
| CMC.BRAIN | LACTB2 | 8 | 71547553 | 71581409 | 0.86 | 3.92E-01 |
| **YFS.BLOOD** | **MS4A6A** | **11** | **59939081** | **59952139** | **7.37** | **1.73E-13** |
| **NTR.BLOOD** | **MS4A6A** | **11** | **59939081** | **59952139** | **6.77** | **1.29E-11** |
| **GTEx7_Whole_Blood** | **MS4A6A** | **11** | **59939081** | **59952139** | **6.55** | **5.93E-11** |
| GTEx7_Whole_Blood | MYBPC3 | 11 | 47352957 | 47374253 | 3.29 | 9.97E-04 |
| YFS.BLOOD | PLIN2 | 9 | 19108373 | 19149288 | -1.51 | 1.30E-01 |
| NTR.BLOOD | PLIN2 | 9 | 19108373 | 19149288 | -2.36 | 1.82E-02 |
| **YFS.BLOOD** | **PTK2B** | **8** | **27168999** | **27316903** | **5.05** | **4.33E-07** |
| **NTR.BLOOD** | **PTK2B** | **8** | **27168999** | **27316903** | **5.90** | **3.74E-09** |
| **GTEx7_Whole_Blood** | **PTK2B** | **8** | **27168999** | **27316903** | **4.95** | **7.30E-07** |
| **GTEx7_Whole_Blood** | **PVR** | **19** | **45147098** | **45166850** | **-3.93** | **8.54E-05** |
| CMC.BRAIN | PVR | 19 | 45147098 | 45166850 | -5.98 | 2.19E-09 |
| YFS.BLOOD | SPI1 | 11 | 47376411 | 47400127 | 3.88 | 1.06E-04 |
| YFS.BLOOD | STX3 | 11 | 59480929 | 59573354 | -0.80 | 4.23E-01 |

**Supplementary Table 7.** Transcriptome-wide association study (TWAS) test statistics for genes in NTR blood, YFS blood and GTEx Whole blood and CMC brain (DLPFC) that were TWAS-significant in monocytes. K_TWAS.Z denotes the gene-level TWAS Z-score and K_TWAS.P denotes the TWAS p-value using the Kunkle 2019 summary statistics (Kunkle *et al.*, 2019). TWAS significant genes after multiple testing correction (Bonferroni) are shown in bold.

**Conditional analysis plots**

Manhattan plots of the Kunkle 2019 GWAS data before (grey) and after (blue) conditioning on the green genes. SNPS are represented by coloured points plotted on the x axis by genomic location. The genes in the locus are represented at the top of the plot. Blue genes are marginally TWAS associated genes and green genes are jointly significant genes. The monocyte cell strain in which genes in the locus were TWAS-significant are shown in each plot.

**1. Locus: PTK2B**

**Monocyte cell strain: CD14, LPS24:**

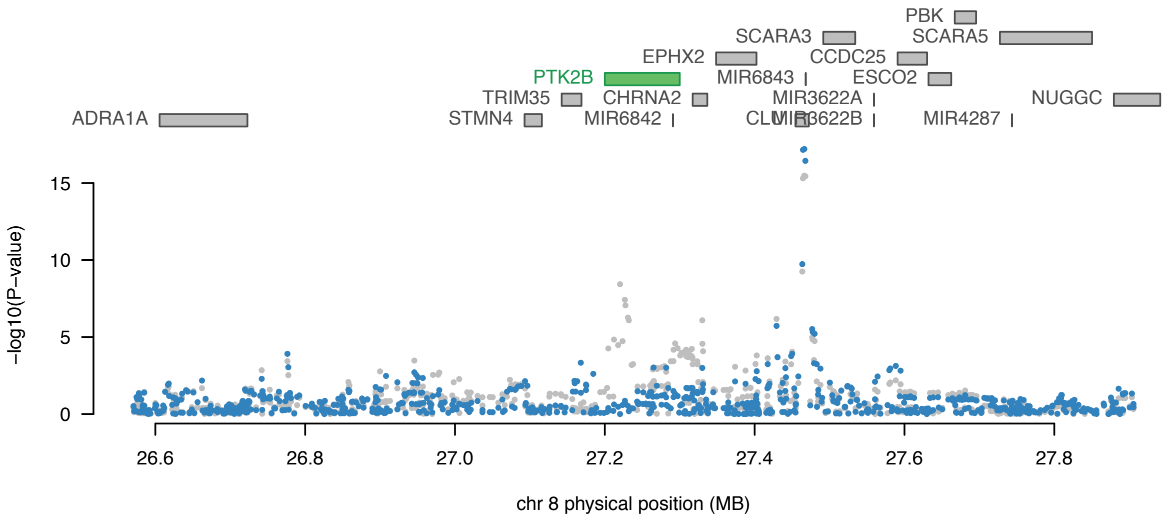

**2. Locus: LACTB2**

**Monocyte cell strain: CD14, IFN, LPS24:**

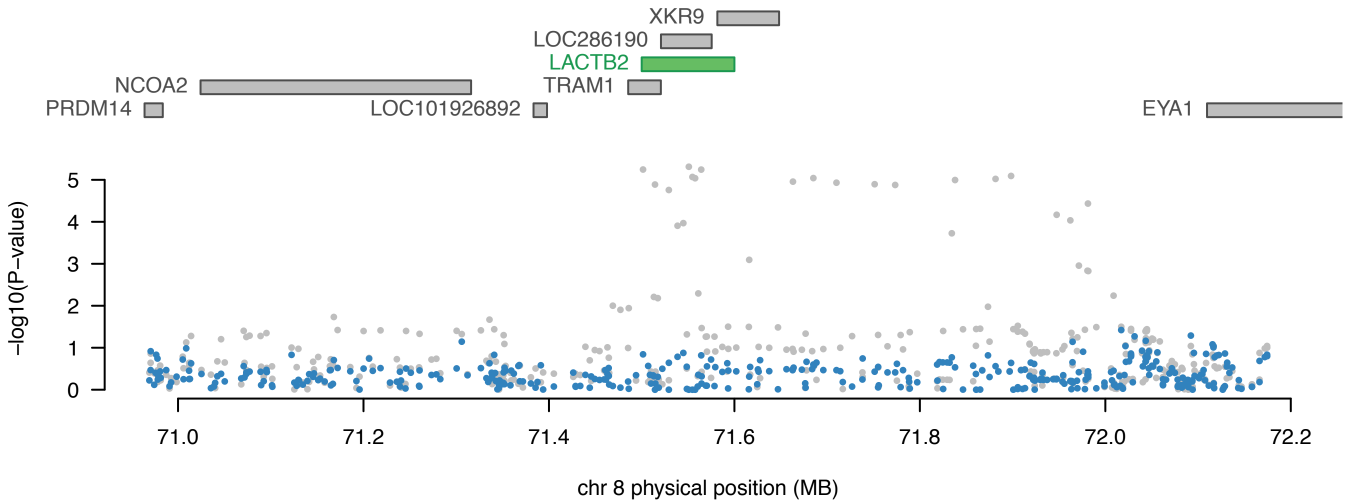

**3. Locus: SPI1**

**Monocyte cell strain: CD14, IFN, LPS2, LPS24:**

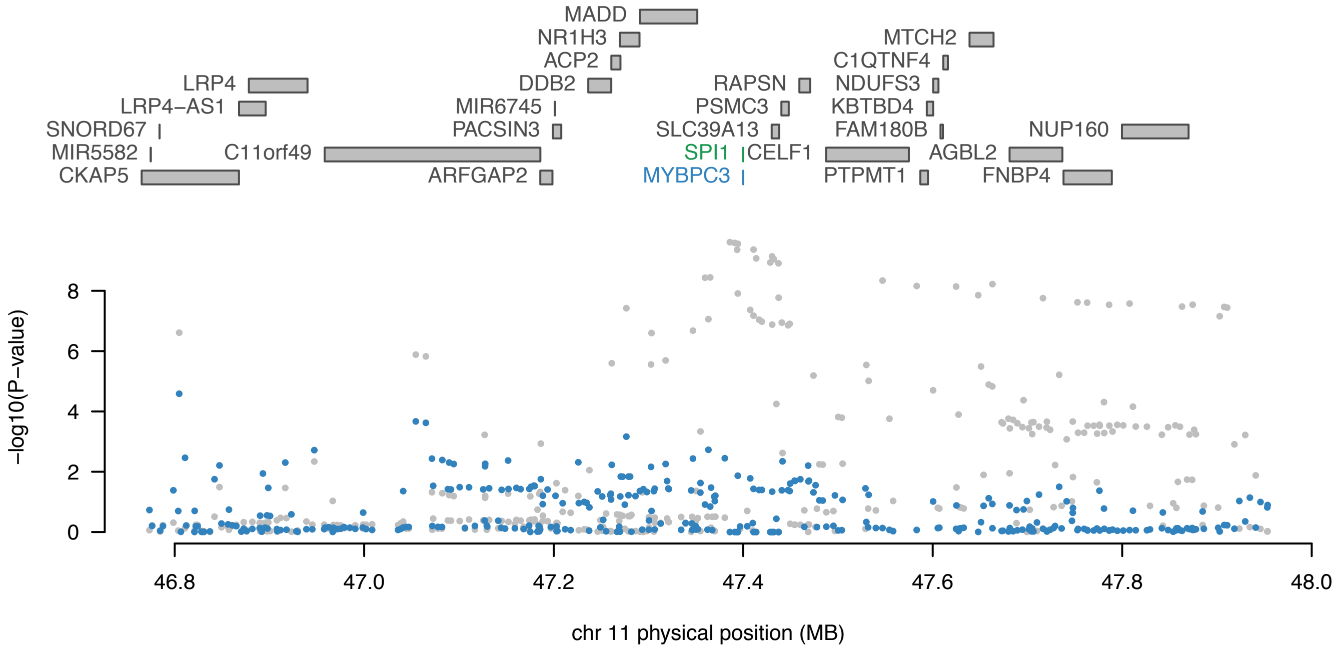

##### 4. Locus: MS4A4A

##### Monocyte cell strain: CD14

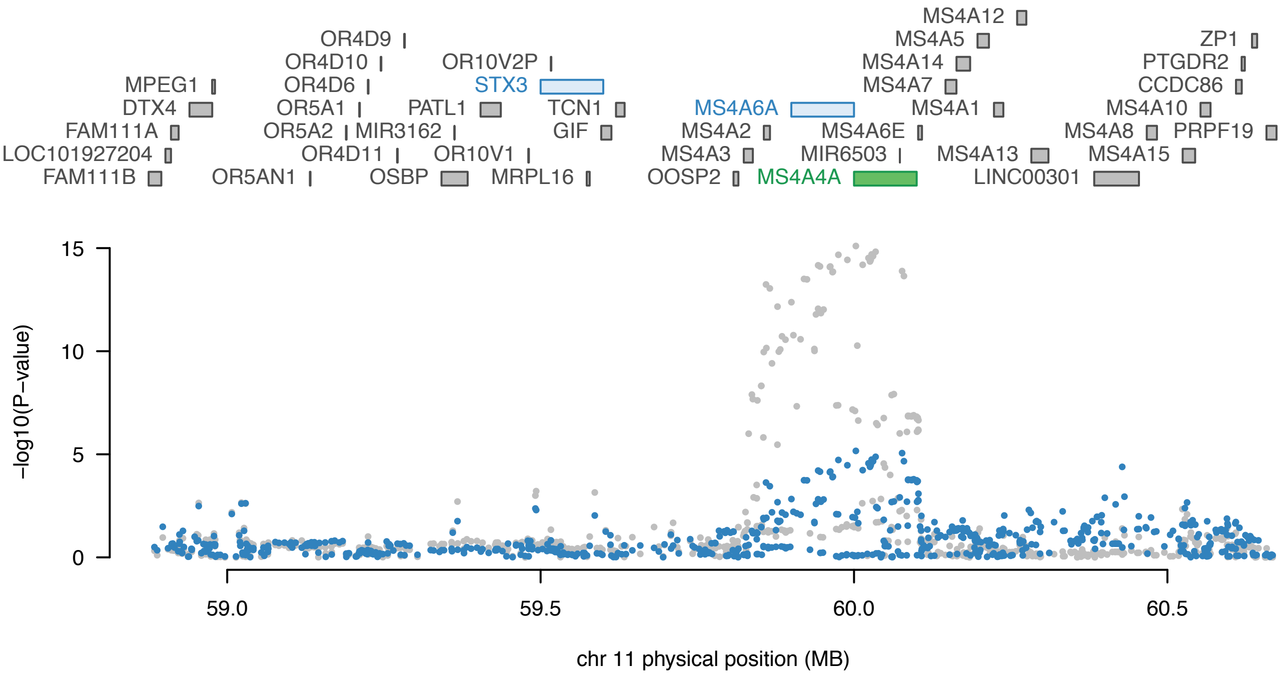

**5.  Locus: BIN1**

**Monocyte cell strain: LPS2, LPS24**

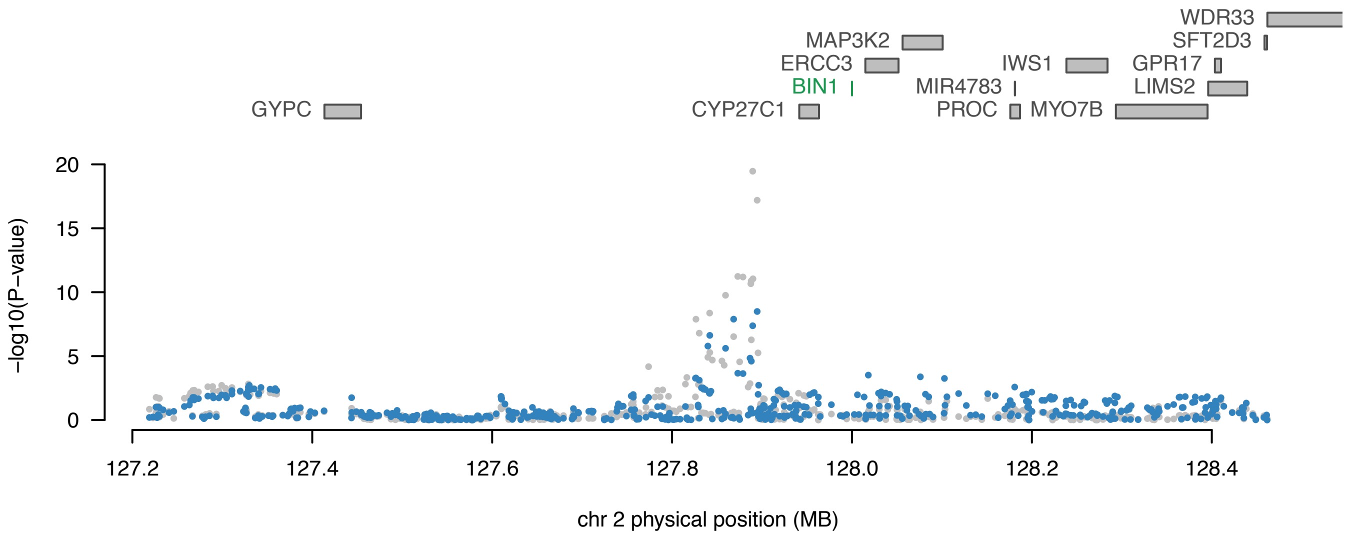

**6. Locus: PLIN1**

**Monocyte cell strain: LPS2**

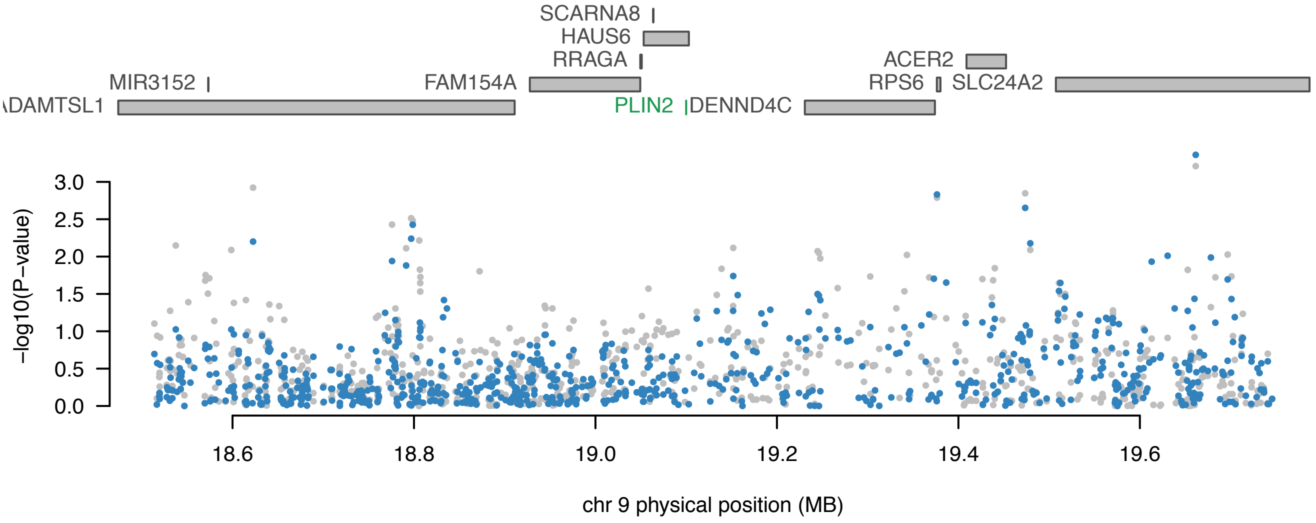

**7.  Monocyte cell strain: LPS2, LPS24**

**Locus: MS4A6E**

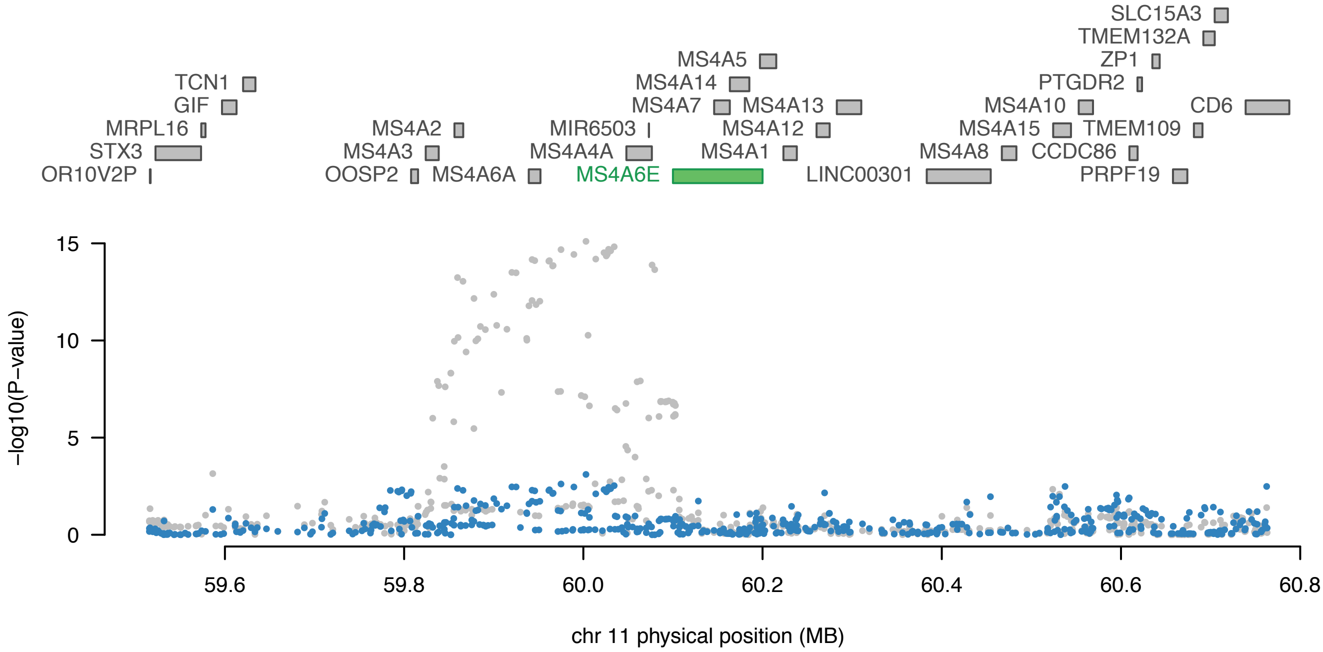

#### 8. Locus: APOE

#### Monocyte cell strain: LPS2

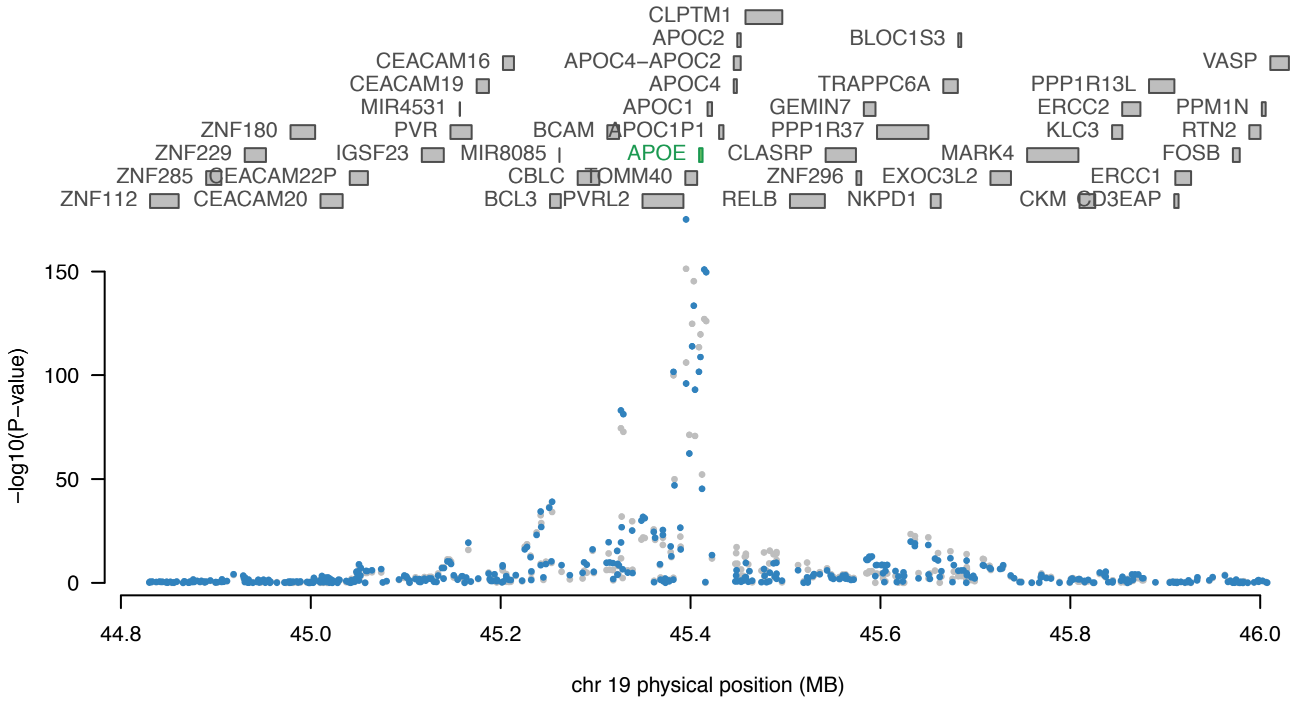

#### 9. Locus: PVR

#### Monocyte cell strain: LPS24

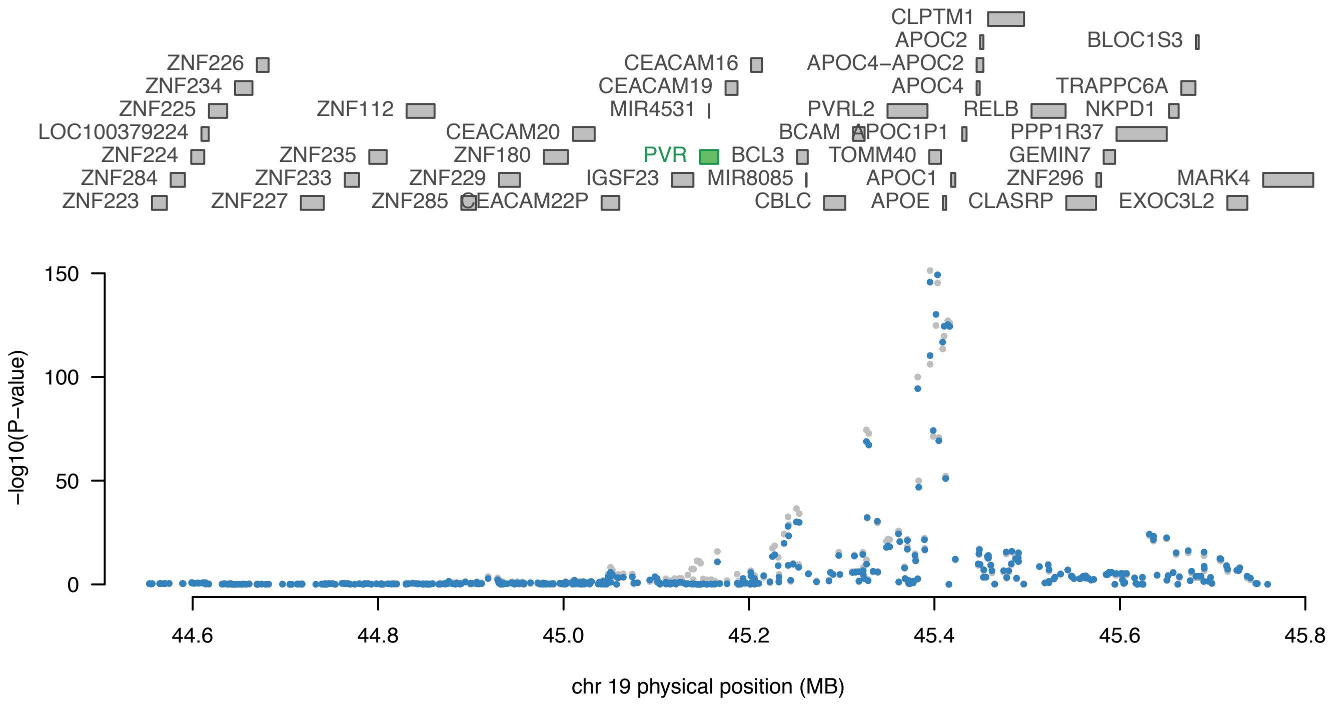

**Supplementary references.**

Bowers JR, Readler JM, Sharma P, Excoffon KJDA. Poliovirus Receptor: More than a simple viral receptor. Virus Res 2017; 242: 1–6.

Corder E, Saunders A, Strittmatter W, Schmechel D, Gaskell P, Small G, et al. Gene dose of apolipoprotein E type 4 allele and the risk of Alzheimer's disease in late onset families. Science (80- ) 1993; 261: 921–923.

Fairfax BP, Makino S, Radhakrishnan J, Plant K, Leslie S, Dilthey A, et al. Genetics of gene expression in primary immune cells identifies cell type-specific master regulators and roles of HLA alleles. Nat Genet 2012; 44: 502–10.

Farrer LA, Cupples LA, Haines JL, Hyman B, Kukull WA, Mayeux R, et al. Effects of age, sex, and ethnicity on the association between apolipoprotein E genotype and Alzheimer disease: A meta-analysis. J Am Med Assoc 1997; 278: 1349–1356.

Hollingworth P, Harold D, Sims R, Gerrish A, Lambert JC, Carrasquillo MM, et al. Common variants at ABCA7, MS4A6A/MS4A4E, EPHA1, CD33 and CD2AP are associated with Alzheimer's disease. Nat Genet 2011; 43: 429–436.

Huang KL, Marcora E, Pimenova AA, Di Narzo AF, Kapoor M, Jin SC, et al. A common haplotype lowers PU.1 expression in myeloid cells and delays onset of Alzheimer's disease. Nat Neurosci 2017; 20: 1052–1061.

Jansen IE, Savage JE, Watanabe K, Bryois J, Williams DM, Steinberg S, et al. Genome-wide meta-analysis identifies new loci and functional pathways influencing Alzheimer's disease risk. Nat Genet 2019; 51: 404–413.

Karch CM, Ezerskiy LA, Bertelsen S, Goate AM, Albert MS, Albin RL, et al. Alzheimer's disease risk polymorphisms regulate gene expression in the ZCWPW1 and the CELF1 loci. PLoS One 2016; 11: e0148717.

Katsumata Y, Nelson PT, Estus S, Fardo DW. Translating Alzheimer's disease–associated polymorphisms into functional candidates: a survey of IGAP genes and SNPs. Neurobiol Aging 2019; 74: 135–146.

Kunkle BW, Grenier-Boley B, Sims R, Bis JC, Damotte V, Naj AC, et al. Genetic meta-analysis of diagnosed Alzheimer's disease identifies new risk loci and implicates Aβ, tau, immunity and lipid processing. Nat Genet 2019 513 2019; 51: 414.

Lambert JC, Ibrahim-Verbaas CA, Harold D, Naj AC, Sims R, Bellenguez C, et al. Meta-analysis of 74,046 individuals identifies 11 new susceptibility loci for Alzheimer's disease.

Nat Genet 2013; 45: 1452–1458.
Lv H, Zhang M, Shang Z, Li J, Zhang S, Lian D, et al. Genome-wide haplotype association study identify the FGFR2 gene as a risk gene for Acute Myeloid Leukemia. Oncotarget 2017; 8: 7891–7899.
Marioni RE, Visscher PM, Harris SE, Gale CR, Yang J, Zhang Q, et al. GWAS on family history of Alzheimer's disease. Transl Psychiatry 2018; 8: 99.
Naj AC, Jun G, Beecham GW, Wang LS, Vardarajan BN, Buros J, et al. Common variants at MS4A4/MS4A6E, CD2AP, CD33 and EPHA1 are associated with late-onset Alzheimer's disease. Nat Genet 2011; 43: 436–443.
Reymond N, Imbert A-M, Devilard E, Fabre S, Chabannon C, Xerri L, et al. DNAM-1 and PVR regulate monocyte migration through endothelial junctions. J Exp Med 2004; 199: 1331–41.
Rosenthal SL, Barmada MM, Wang X, Demirci FY, Kamboh MI. Connecting the dots: Potential of data integration to identify regulatory snps in late-onset alzheimer's disease GWAS findings. PLoS One 2014; 9: e95152.
Seshadri S, Fitzpatrick AL, Ikram MA, DeStefano AL, Gudnason V, Boada M, et al. Genome-wide analysis of genetic loci associated with Alzheimer disease. JAMA - J Am Med Assoc 2010; 303: 1832–1840.
Zeller T, Wild P, Szymczak S, Rotival M, Schillert A, Castagne R, et al. Genetics and beyond - the transcriptome of human monocytes and disease susceptibility. PLoS One 2010; 5: e10693.
Zhernakova D V., Deelen P, Vermaat M, Van Iterson M, Van Galen M, Arindrarto W, et al. Identification of context-dependent expression quantitative trait loci in whole blood. Nat Genet 2017; 49: 139–145.

URLs

1. https://www.ebi.ac.uk/arrayexpress/experiments/E-MTAB-2232/.
2. Fairfax *et al* Supplementary Material: https://science.sciencemag.org/content/suppl/2014/03/05/343.6175.1246949.DC1?_ga=2.250430726.142617004.1580743814-1392779675.1580743814
3. Illumina HumanHT-12_V4_0_R1_15002873_B array: https://www.ebi.ac.uk/arrayexpress/files/A-MEXP-2210/A-MEXP-2210.adf.txt
4. Biomart: https://bioconductor.org/packages/release/bioc/html/biomaRt.html
5. WGCNA: https://horvath.genetics.ucla.edu/html/CoexpressionNetwork/Rpackages/WGCNA/
6. Liftover tool: https://genome.ucsc.edu/cgi-bin/hgLiftOver
7. Plink 1.9 http://www.cog-genomics.org/plink/1.9/
8. LD Score Regression (LDSC) software : v1.0.0 : https://github.com/bulik/ldsc
9. HapMap 3 https://www.sanger.ac.uk/resources/downloads/human/hapmap3.html
10. Oliver Pain: OP_packaging_fusion_weights.R : https://github.com/opain/Calculating-FUSION-TWAS-weights-pipeline.
11. FUSION software http://gusevlab.org/projects/fusion/
12. UpSetR: https://cran.r-project.org/web/packages/UpSetR
13. Common mind consortium: https://www.nimhgenetics.org/resources/commonmind
